# Neural evidence for a two-stage model of visual processing

**DOI:** 10.1101/2024.11.15.623853

**Authors:** Maëlan Q. Menétrey, Michael H. Herzog, David Pascucci

## Abstract

In some forms of postdictive phenomena, later events influence the perception of earlier ones, suggesting that conscious perception may be preceded by extended periods of unconscious processing. An example is the Sequential Metacontrast (SQM) paradigm, in which vernier offsets are unconsciously integrated over several hundred milliseconds before conscious perception. Obviously, the integrated percept can only emerge after each individual element in the stream has been processed. Thus, the SQM provides a unique opportunity to dissociate unconscious from conscious stages of visual processing, as these stages are well separated in time. Using EEG during the SQM, we identified two distinct stages of neural activity: an early occipital EEG activity pattern (∼200 ms after the initial vernier) associated with unconscious processing, and a later centro-parietal EEG pattern (∼400-600 ms after SQM onset) associated with the integrated percept and the behavioral report. We propose that the transition between these neural patterns marks the shift from unconscious encoding of individual visual stimuli to their integrated percept.

## Introduction

Perception feels like a continuous stream: we seem to consciously experience the world at each moment of time. However, several visual phenomena challenge this idea. For instance, in a classic apparent motion phenomenon, two dots of different colors are flashed at separate locations with a brief delay (Kolers & Von Grünau, 1976). Rather than perceiving two distinct flashes one after the other, we perceive a single moving dot that changes color midway. Logically, the color change can only be perceived after the second dot is presented, implying that later events influence how we perceive earlier ones. These postdictive effects provide a unique opportunity to study the neural correlates of both conscious and unconscious processing, as well as the transitions between the two.

A prime tool for studying these processes is the Sequential Metacontrast (SQM) paradigm (Figures 1A and 1B). In this paradigm, a sequence of vertical lines is presented, creating the illusion of two diverging motion streams expanding from a central point (Otto et al., 2009, 2010). When one of the lines contains a horizontal vernier offset (i.e., the lower segment of the line is slightly shifted leftward or rightward relative to the upper segment), all the other lines are perceived as having the same offset, even though they are actually straight. When the stream contains two verniers with opposite offsets, the offsets integrate: participants are unable to report the individual offsets separately and cannot locate which lines carry offsets (Drissi-Daoudi et al., 2019). Thus, the multiple offsets are integrated unconsciously and the result of the integration is the final conscious percept.

**Figure 1.**
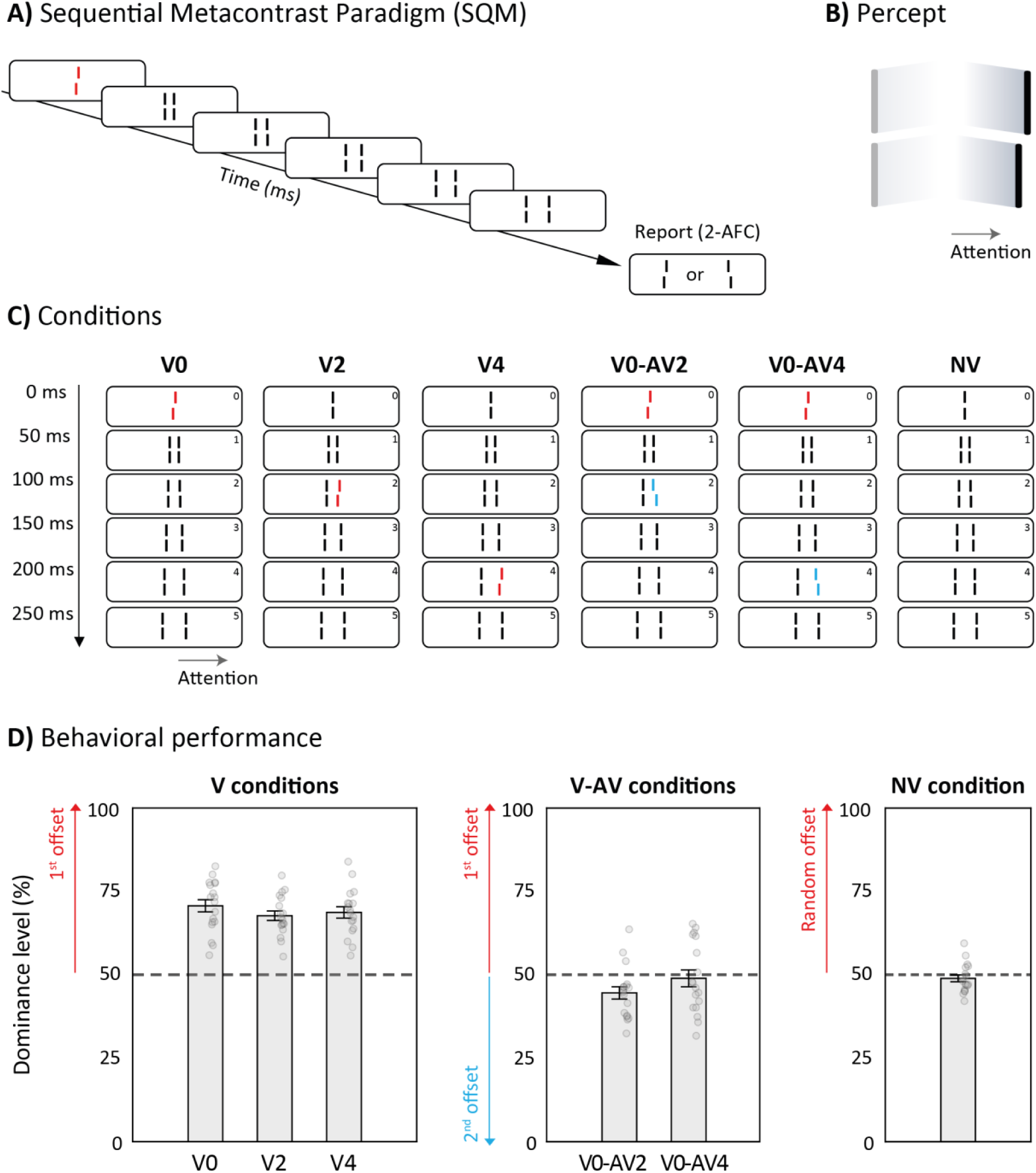
A) The Sequential Metacontrast paradigm (SQM). A central line, here with a left vernier offset, is followed by pairs of flanking lines eliciting a percept of two diverging streams. Participants attend to one stream (in this study, the rightward stream) and, at the end of each trial, report the perceived offset using a hand-held push-button response (left vs right, two-alternative forced choice). B) In the attended stream, the offset is typically perceived as propagating along the entire stream. C) In V conditions, only one vernier is presented, either in the central line (V0), in the second (V2), or in the fourth (V4) flanking line. In V-AV conditions, the central line is offset, and one of the flanking lines is offset in the opposite direction (anti-vernier; V0-AV2 or V0-AV4). In a control condition (NV), only straight lines are shown. The red and blue vernier offsets in the figure are for illustration purposes only; in the experiment, all lines were the same color. D) In V conditions, participants could well report the offset direction. In V-AV conditions, participants could not perceive the individual vernier offsets, and hence performance is at 50%. This indicates that both vernier offsets contribute equally to the integrated percept, or that each offset is reported with equal probability in a given trial. When no vernier was presented or perceived, participants were guessing. In NV condition, performance is determined by comparing the response to a randomly chosen offset.

In the SQM, the integration period of the vernier offsets is approximately 290-450 ms, depending on the participant and the condition (Vogelsang et al., 2023, 2024). When two offsets are separated by more than ∼ 450 ms, participants can report the offsets individually (Drissi-Daoudi et al., 2019; Vogelsang et al., 2023). These findings suggest that the brain integrates features within a relatively long-lasting window of unconscious processing, followed by an integrated conscious percept (Herzog et al., 2016, 2020). The existence of such long-lasting integration windows has been demonstrated not only in visual perception (Scharnowski et al., 2009; Mudrik et al., 2011; Akyürek & Wolff, 2016; Marti & Dehaene, 2017; Sergent, 2018), but also in auditory perception (McWalter & McDermott, 2019) and across different sensory modalities (Fiebelkorn et al., 2010; Stiles et al., 2018; Rimsky-Robert et al., 2019).

In this study, we used electroencephalography (EEG) decoding to investigate the neural mechanisms involved in both the unconscious integration of vernier offsets and the subsequent emergence of a conscious percept. Our approach was guided by the following rationale: in the SQM, regardless of the number and spatiotemporal locations of the verniers, all offsets are *integrated unconsciously*, and only the outcome is *consciously perceived* (Herzog et al., 2020). Neural activity encoding information about the number or location of individual offsets should therefore reflect unconscious processing, whereas neural activity encoding the integrated outcome should reflect processes linked to the content of the conscious percept.

## Results

### Unconscious feature integration within the SQM stream

In the conditions with only a single vernier in the stream (V conditions; Figure 1A and 1C), offset discrimination was above 50% (Figure 1D, left panel). This effect was consistent regardless of where the offset appeared in the stream: whether the offset occurred in the central line or in the second or fourth flanking lines, mean accuracy remained high (70% ± 7%). Paired t-tests comparing each of these V conditions with a no vernier (NV) control condition resulted all in significant differences (all *p* < .001, Cohen’s *d’* > 3.3; Figure 1D, left panel).

In contrast, when two opposite vernier offsets (a vernier and an anti-vernier) were included in the stream (V-AV condition, Figure 1C), mean accuracy was 47% ± 9%, comparable to the NV condition (paired t-tests between the V-AV and NV conditions: all *p* > .05; Figure 1D, right panel). Thus, conscious perception does not depend on the spatiotemporal location and number of offsets in the stream, as there is no conscious percept of the individual elements.

### Neural representations preserve the chronology of events in the SQM stream

We first aimed to decode neural activity related to the processing of individual vernier offsets. To this end, we applied Linear Discriminant Analysis (LDA) to EEG activity patterns to decode the presence and spatiotemporal location of vernier offsets across the stream.

We ran separate LDA classifiers to discriminate between trials with no offset (NV condition) and trials with a single vernier offset presented at different spatiotemporal locations within the stream (V0: 0 ms, V2: 100 ms, V4: 200 ms, Figure 1C). Using a temporal generalization approach, the classifiers were trained on data from each time point and tested across all other time points (see Methods). The classifiers successfully discriminated the presence of the vernier offsets in all V conditions (Area Under the Curve, AUC > 0.5, one-tailed cluster-based permutation test, *p* < .05; Figures 2A and 2B). The resulting patterns were similar across conditions with decoders generalizing only over short time windows, as indicated by higher decoder performance when both training and testing occurred at the same time points (i.e., the diagonal elements of the temporal generalization matrices).

**Figure 2.**
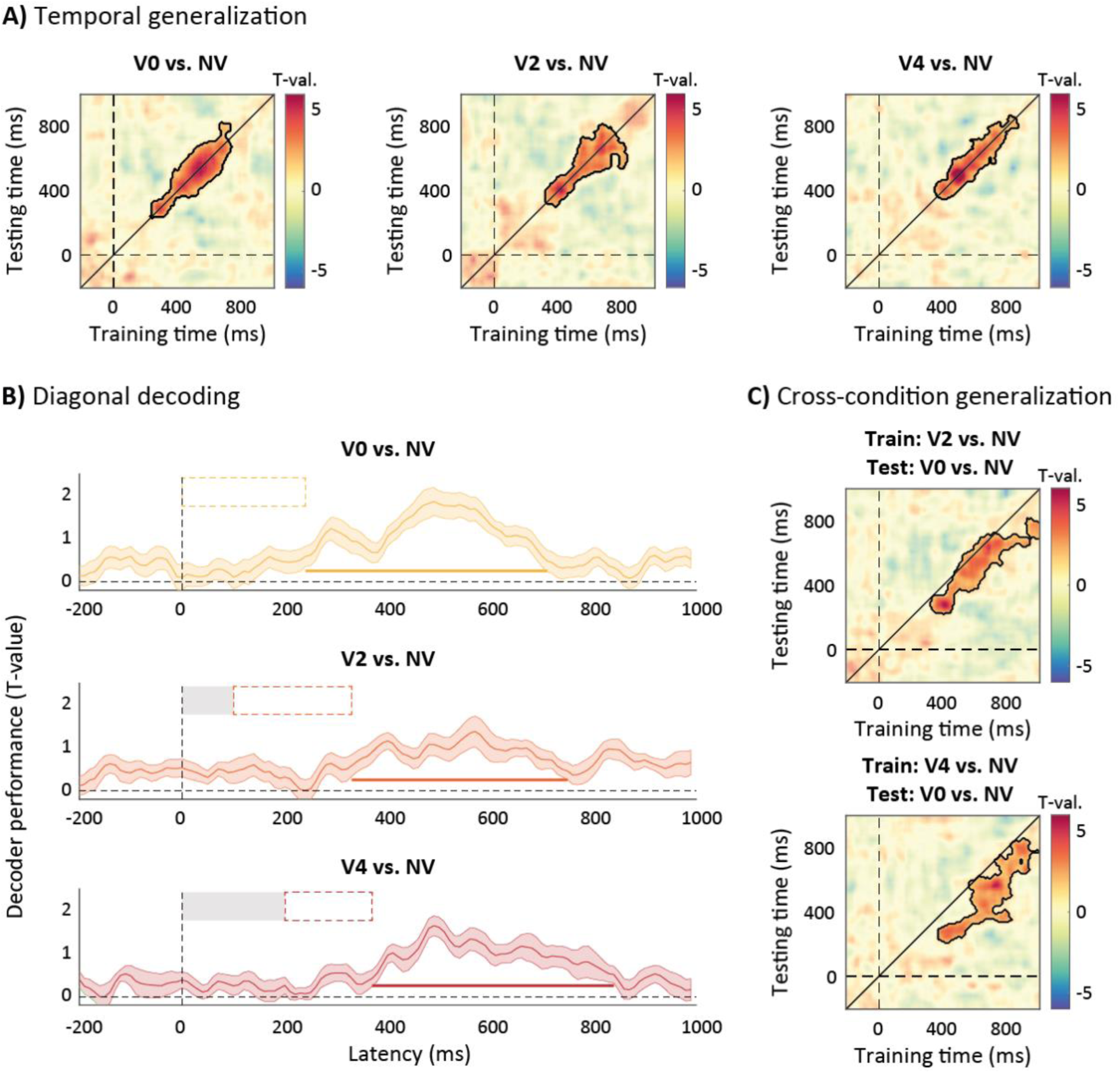
Linear discriminant analysis (LDA) with temporal and cross-condition generalization to decode conditions with one vernier and no vernier. A) Temporal generalization matrices, obtained by training a classifier on data at every time point and testing it at all other time points. The classifiers can successfully discriminate between the conditions (V0, V2, and V4 vs. NV). Significant clusters are highlighted (AUC > 0.5, one-tailed cluster-based permutation test, *p* < .05). B) Diagonal elements of the temporal generalization matrices, representing decoding results when training and testing at the same time point (group average AUC and SEM). Significant time windows are highlighted by the horizontal lines at the bottom. Temporal delays in the decoding results, relative to the actual onset of the vernier in the stream, are indicated by the dashed rectangles. C) Cross-condition generalization matrices showing that classifiers trained with V2 or V4 vs. NV successfully discriminate V0 vs. NV but with earlier onsets corresponding to the effective stimulus delay between V0 and V2 (i.e., 100 ms later) and V4 (i.e., 200 ms later). Significant clusters are highlighted (AUC > 0.5, one-tailed cluster-based permutation test, *p* < .05).

Crucially, the decoding latencies revealed systematic delays that corresponded to the actual spatiotemporal location of the vernier offset in the stream—decoding was successful at 240 ms for V0 vs. NV, at 330 ms for V2 vs. NV, and at 370 ms for V4 vs. NV (Figure 2B). Hence, although participants perceived the offset in the entire stream, the offset was decoded only around 200 ms (170–240 ms) after its physical onset, with slightly decreasing delays when the vernier was presented later in the stream.

This result was further supported by cross-condition decoding, where we trained a classifier on discriminating V2 vs. NV or V4 vs. NV and tested it on V0 vs. NV. This revealed shared neural representations of the single vernier across conditions but shifted along the diagonal in the cross-condition temporal generalization matrix (Figure 2C). That is, a decoder trained to detect the vernier in the V2 or V4 condition successfully decoded it in V0 (AUC > 0.5, one-tailed cluster-based permutation test, *p* < .05), but with earlier onsets of approximately 100 and 200 ms, respectively, corresponding to the actual latencies between conditions.

Decoding the conditions with a single vernier against the condition with no vernier was significant in windows ranging from 240 to 710 ms in V0 vs. NV, 330 to 750 ms in V2 vs. NV, and 370 to 840 ms in V4 vs. NV). In these time windows, we found two distinct EEG activation patterns, i.e., topographies of EEG activity associated with the decoder results (see Methods and Figure 3A). These scalp patterns and their transitions were consistent across participants (Figure 3A, bottom row, Cohen’s *d* of the scalp activation patterns at the group level).

**Figure 3.**
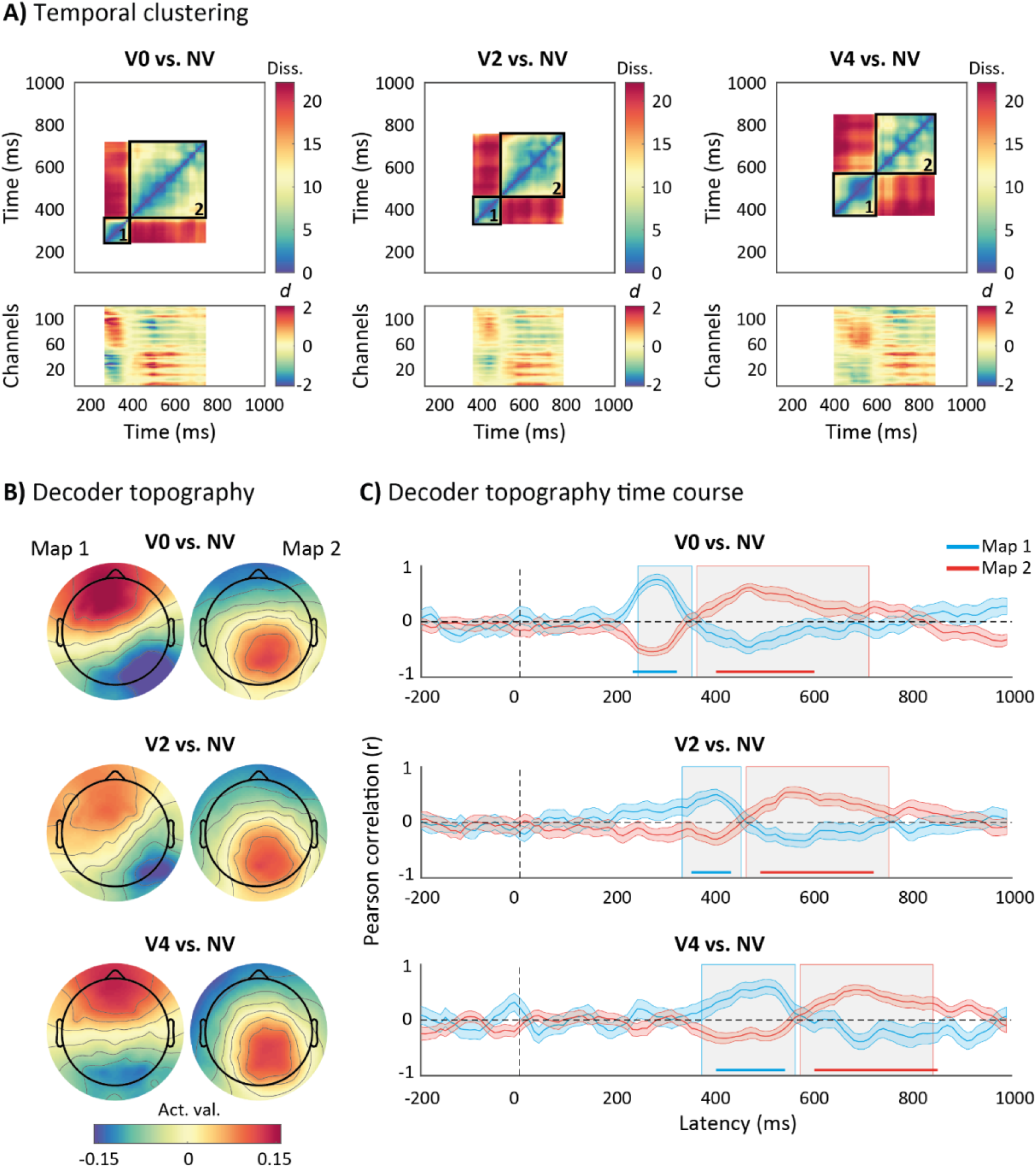
EEG activation patterns contributing to decoding conditions with one vernier and no vernier. A) Analysis of dissimilarity matrices between the EEG activation patterns extracted from significant time points (diagonal elements of the temporal generalization matrices for V0, V2, or V4 vs. NV; see Figure 2B) reveals a clear transition between two main activation patterns (outlined by black squares) for each decoding analysis (upper row). These activations patterns were consistently expressed across participants, with many channels and time points showing large effect sizes (difference between each activation pattern and zero, calculated for each electrode and time point |Cohen’s d| > 1; bottom row). B) Decoder topographies, averaged across participants, are derived from the two main activations patterns identified with the analysis of dissimilarity matrices. Each decoding analysis shows similar topographical maps but with variations in their transition dynamics. C) Temporal correlations between the two identified topographies and the back-projected time course of EEG activation patterns for each decoding analysis. Gray areas highlight the significant decoding windows found in 2B and 3A. Blue and red lines represent the occipital and parietal topographies, respectively, and the shaded areas indicate SEM. Significant positive correlations are also highlighted in blue for the occipital topography or in red for the parietal topography (Pearson’s *r* > 0, one-tailed cluster-based permutation test, *p* < .05).

The first topography displayed a stronger contribution of occipital electrodes (hereafter referred to as the “occipital topography”), with a duration that increased as the vernier offset occurred later in the stream (Figure 3B). In contrast, the second topography showed a stronger contribution of parietal electrodes (hereafter referred to as the “parietal topography”), with a duration that decreased as the vernier offset occurred later (Figure 3B). The time course of these topographies, estimated by back-projecting activation patterns onto the original EEG time series (see Methods), revealed a clear transition between the two patterns, with evident temporal shifts across conditions (i.e., the transition occurred around 360 ms in V0 vs. NV, 460 ms in V2 vs. NV, and 570 ms in V4 vs. NV; Figure 3C). We further confirmed the robustness of the effect across participants by estimating the proportion of individual activation patterns showing a positive correlation with the occipital topography within the time window corresponding to the first cluster (17/18 participants in V0 vs. NV, 15/18 in V2 vs. NV, and 15/18 in V4 vs. NV), as well as those showing a positive correlation with the parietal topography within the time window corresponding to the second cluster (17/18 participants in V0 vs. NV, 15/18 in V2 vs. NV, and 17/18 in V4 vs. NV).

### Neural correlates of unconscious processing

Up to this point, we have focused on conditions featuring a single vernier, whose offset is consciously perceived throughout the entire stream. Even though the spatiotemporal location of the vernier was not consciously perceived, it could still be decoded via two distinct EEG topographies. Next, we analyzed conditions involving two opposite vernier offsets, separated by varying intervals (V0-AV2: 50 ms, V0-AV4: 150 ms; Figure 1C), using LDA to distinguish these conditions from a condition featuring only straight lines (NV).

In the V-AV conditions, behavioral performance was around 50% (Figure 1D). As previously suggested, this may reflect either that both offsets are fully integrated, resulting in the perception of a straight line as in the NV condition (Drissi-Daoudi et al., 2019), or that there is an equal probability that one of the two vernier offsets “wins” the integration process in a given trial (Menétrey et al., 2023).

The decoder successfully discriminated V0-AV2 or V0-AV4 trials from NV trials (AUC > 0.5, one-tailed cluster-based permutation test, *p* < .05; Figure 4A). In both analyses, the significant decoding windows (170-1000 ms; Figure 4B) exhibited a similar sequence of EEG activation patterns (Figure 4C), with topographies (Figure 4D) closely resembling those observed in the V0 vs. NV decoding analysis (Figure 3B). Similarly, the significant decoding onsets and the transition dynamics around 370 ms (Figures 4B and 4E) closely mirrored those found in V0 vs. NV conditions (Figures 2B and 3C for comparison). The effect was also robust across participants (positive correlation with the occipital topography within the time window associated with the first cluster: 18/18 participants in V0-AV2 vs. NV and 14/18 in V0-AV4 vs. NV; positive correlation with the parietal topography within the time window associated with the second cluster: 16/18 participants in V0-AV2 vs. NV and 18/18 in V0-AV4 vs. NV). Thus, EEG activation patterns distinguished the presence of two offsets from no offsets, even though the individual offsets were integrated and not perceived separately.

**Figure 4.**
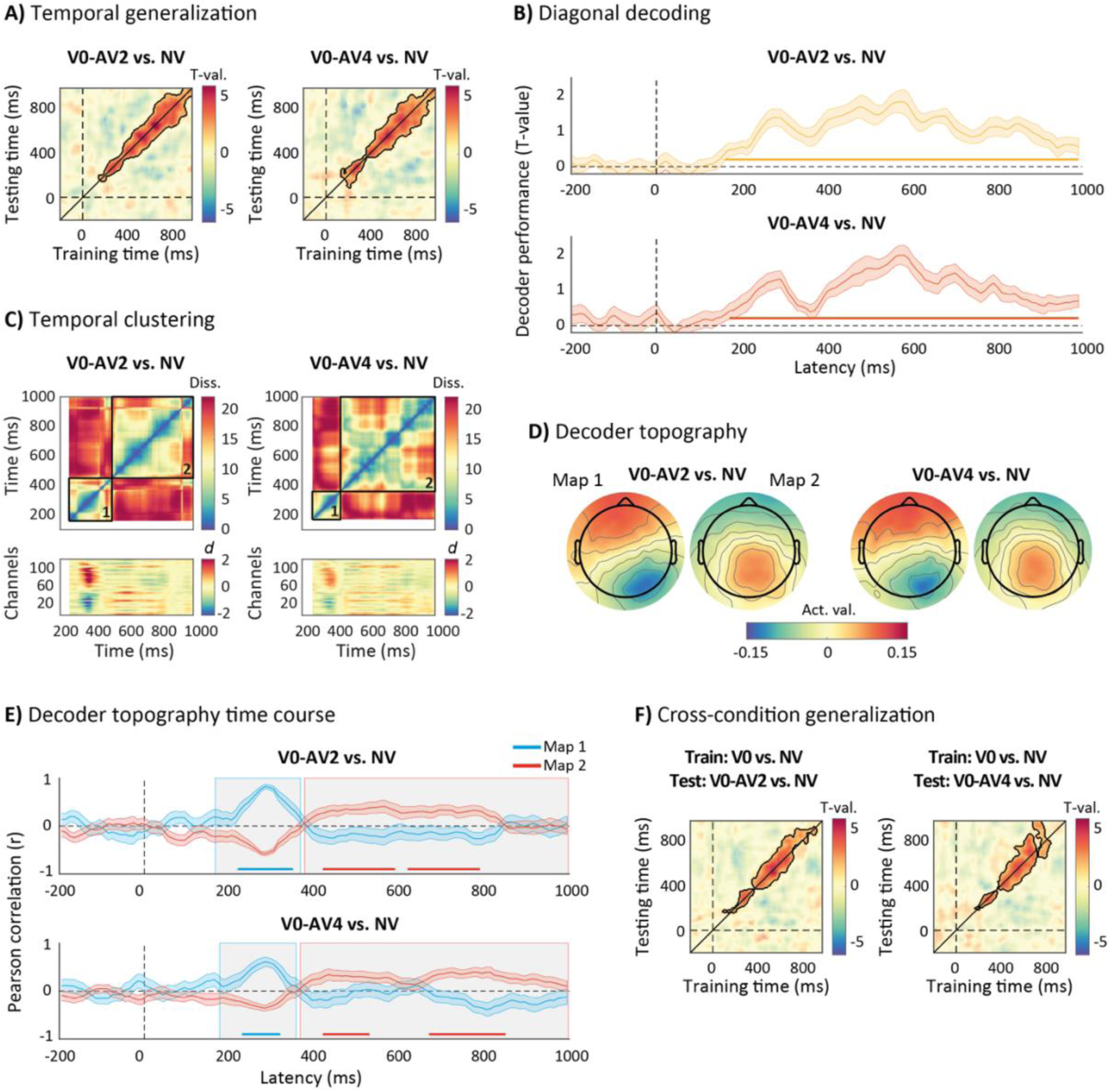
LDA with temporal and cross-condition generalization to decode conditions with two opposite verniers and no vernier. A) Temporal generalization matrices showing that classifiers successfully discriminate the conditions (V0-AV2 or V0-AV4 vs. NV). Significant clusters are highlighted (AUC > 0.5, one-tailed cluster-based permutation test, *p* < .05). B) Diagonal elements of the temporal generalization matrices for V0-AV2 or V0-AV4 vs. NV, representing decoding results when training and testing at the same time point (group average AUC and SEM). Significant time windows are highlighted by the horizontal lines at the bottom. C) Analysis of dissimilarity matrices between the EEG activation patterns extracted from significant time points reveals a clear transition between two main activation patterns (outlined by black squares) for each decoding analysis (upper row). These activations patterns were consistently expressed across participants, with many channels and time points showing large effect sizes (difference between each activation pattern and zero, calculated for each electrode and time point |Cohen’s d| > 1; bottom row). D) Decoder topographies, averaged across participants, are derived from the two main activation patterns identified with the analysis of dissimilarity matrices. Each decoding analysis shows similar topographical maps and transition dynamics. E) Temporal correlation between the two identified topographies and the back-projected time course of EEG activation patterns for each decoding analysis. Gray areas highlight the significant decoding windows found in 4B and 4C. Blue and red lines represent the occipital and parietal topographies, respectively, and the shaded areas indicate SEM. Significant positive correlations are also highlighted in blue for the occipital map or in red for the parietal (Pearson’s *r* > 0, one-tailed cluster-based permutation test, *p* < .05). F). Cross-condition generalization matrices showing that classifiers trained with V0 vs. NV successfully discriminate V0-AV2 or V0-AV4 vs. NV with similar onset. Significant clusters are highlighted (AUC above 0.5, one-tailed cluster-based permutation test, *p* < .05).

Furthermore, cross-condition decoding revealed that a decoder trained to detect the vernier in the V0 condition (vs. NV) still successfully decoded, with similar onset and temporal dynamics, V0-AV2 or V0-AV4 conditions from NV condition (AUC > 0.5, one-tailed cluster-based permutation test, *p* < .05, Figure 4F), indicating shared neural representations between conditions with a single vernier and two opposite vernier offsets.

Importantly, we also found that the two V-AV conditions, where the second vernier occurred at different times (V0-AV2 vs. V0-AV4), could be decoded from V0 trials, where only a single vernier was shown and consciously perceived (AUC > 0.5, one-tailed cluster-based permutation test, *p* < .05; Figure 5A). The corresponding decoding windows (from 300 ms for V0-AV2 vs. V0, and from 340 ms for V0-AV4 vs. V0) were compatible with the latencies observed when decoding conditions in which a single vernier was physically presented later in the stream (V2 or V4 vs NV, Figure 2B). Similarly, the two V-AV conditions were also reliably discriminated from each other (AUC > 0.5, one-tailed cluster-based permutation test, *p* < .05; Figure 5A). That is, EEG activation patterns contained information about the distinct spatiotemporal location of the second vernier, which was decodable starting from 420 ms onward. These results suggest that neural representations of the second vernier persisted despite the second offset not being typically consciously perceived as a distinct event at its specific spatiotemporal location (Otto, 2006; Drissi-Daoudi et al., 2019).

**Figure 5.**
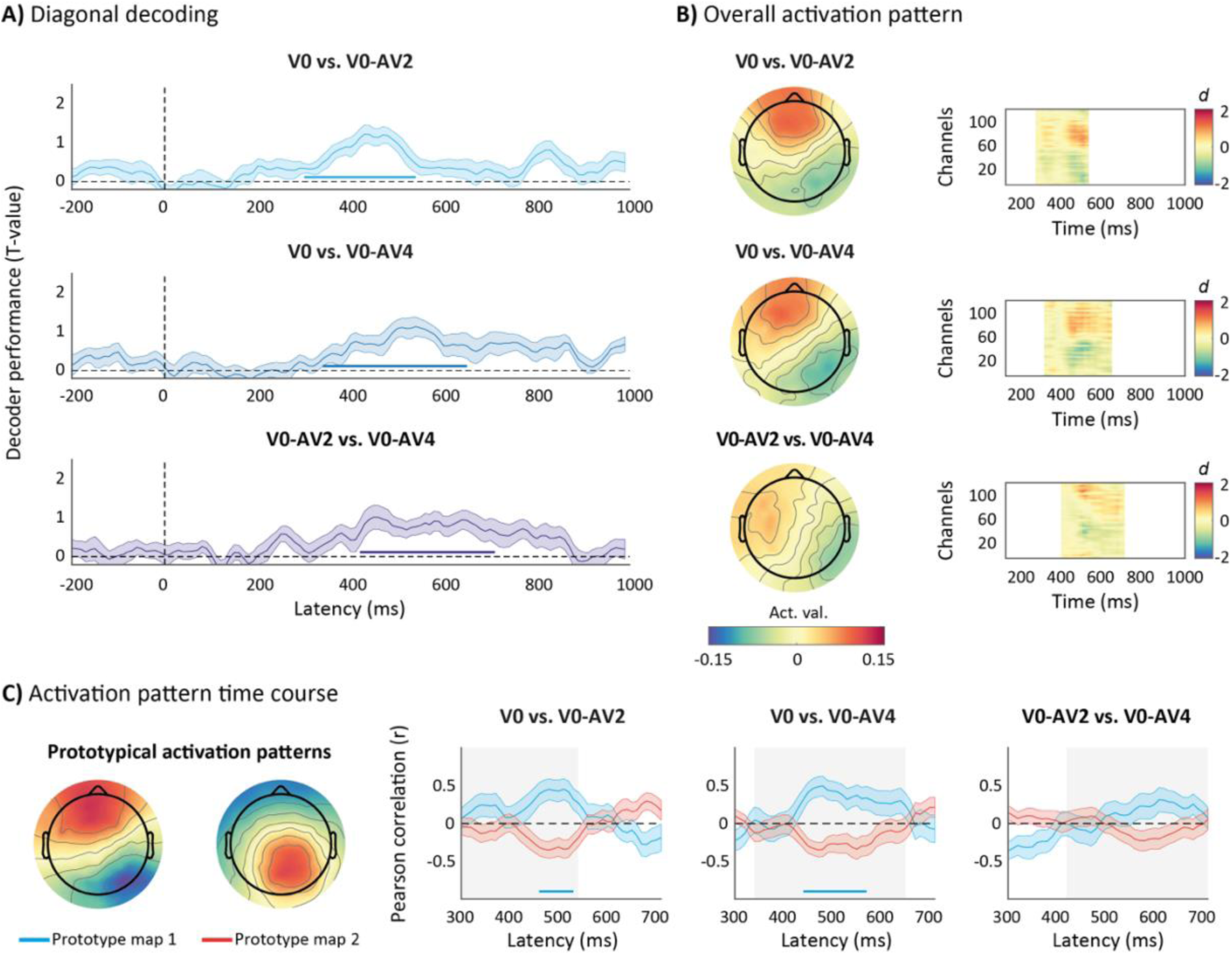
LDA between conditions with one and two verniers and between conditions with two verniers. A) Diagonal elements of the temporal generalization matrices (shown in Supplementary Figure 2), representing decoding results when training and testing at the same time point (group average AUC and SEM). Significant time windows are highlighted by the horizontal lines at the bottom, showing that classifiers successfully discriminate the conditions (V0 vs. V0-AV2 or V0-AV4, and V0-AV2 vs. V0-AV4; AUC > 0.5, one-tailed cluster-based permutation test, *p* < .05). B) The decoder topographies, averaged across participants, are derived from the averaged activation pattern over the entire significant window of each decoding analysis. These activations patterns were consistently expressed across participants, with many channels and time points showing large effect sizes (difference between each activation pattern and zero, calculated for each electrode and time point |Cohen’s d| > 1; bottom row). C) Prototypical activation patterns (left side) reflecting the occipital (map 1) and parietal topographies (map 2), corresponding to the average of the two distinct activation patterns identified via dissimilarity matrix analysis for all V or V-AV vs. NV contrasts (see Figures 3B and 4D), were temporally correlated with the activation patterns found in each decoding window (right side). Gray areas highlight the significant decoding window found in 5A. Blue and red lines represent the occipital and parietal topographies, respectively, and the shaded areas indicate SEM. Significant positive correlations are also highlighted in blue for the occipital topography or in red for the parietal topography (Pearson’s *r* > 0, one-tailed cluster-based permutation test, *p* < .05).

The average activation pattern contributing to these decoding results (Figure 5B) resembled the occipital topography identified when contrasting conditions with one or two offsets from the condition without any offset (Figures 3B and 4D). This topography dominated and remained stable throughout the significant decoding windows and across the three analyses (V0 vs. V-AV and V0-AV2 vs. V0-AV4; see Figure 5B, effect size plots). To further validate this finding, we correlated the activation patterns obtained here with the two prototypical topographies derived from averaging across all V or V-AV vs. NV decoding analyses (Figures 3B and 4D; see Methods). The results confirmed the dominant contribution of the occipital prototypical topography (Figure 5C), temporally aligned with the onset of the second vernier. We further corroborated the dominant occipital contribution by decoding between V conditions involving physically different sequences of events that nevertheless led to the common conscious perception of a single integrated offset (V0 vs. V2 or V4, and V2 vs. V4). These results further suggest that this topography reflects the processing of the spatiotemporal location of a vernier offset prior to its integration within the SQM stream (see Supplementary Figure 1).

Based on these findings, we interpret the occipital topography as the neural correlate of unconscious processing of the vernier stimuli, occurring prior to the formation of the integrated percept. Conversely, the presence of the parietal topography when discriminating V-AV from NV trials (Figure 4D) might reflect the content of the final integrated percept. Indeed, the latencies and sequence of topographies in these decoding analyses are similar to those observed in the condition with a single vernier presented in the central line (V0; Figure 3B), consistent with the view that, in V-AV conditions, participants may still perceive a residual offset in the stream (Menétrey et al., 2023). Such a percept could arise from a non-uniform integration of the first and second vernier offsets, resulting in one offset contributing more strongly than the other (see also Supplementary Figure 3 for additional cross-condition generalization evidence between V and V-AV conditions, as a function of whether the first or second vernier dominated the reported percept).

### Neural correlates of the integrated percept

To directly assess the involvement of the parietal topography in conscious processing, we applied LDA to decode correct vs. incorrect reports of the offset direction in the conditions featuring only a single vernier (V conditions). As shown in Figure 6, the classifiers successfully decoded the performance in all single-vernier conditions (Figure 6A; AUC > 0.5, one-tailed cluster-based permutation test, *p* < .05). However, significant decoding emerged at later latencies—480 ms for V0, 460 ms for V2, and 570 ms for V4— compared to the earlier latencies observed when decoding V or V-AV conditions against the NV condition.

**Figure 6.**
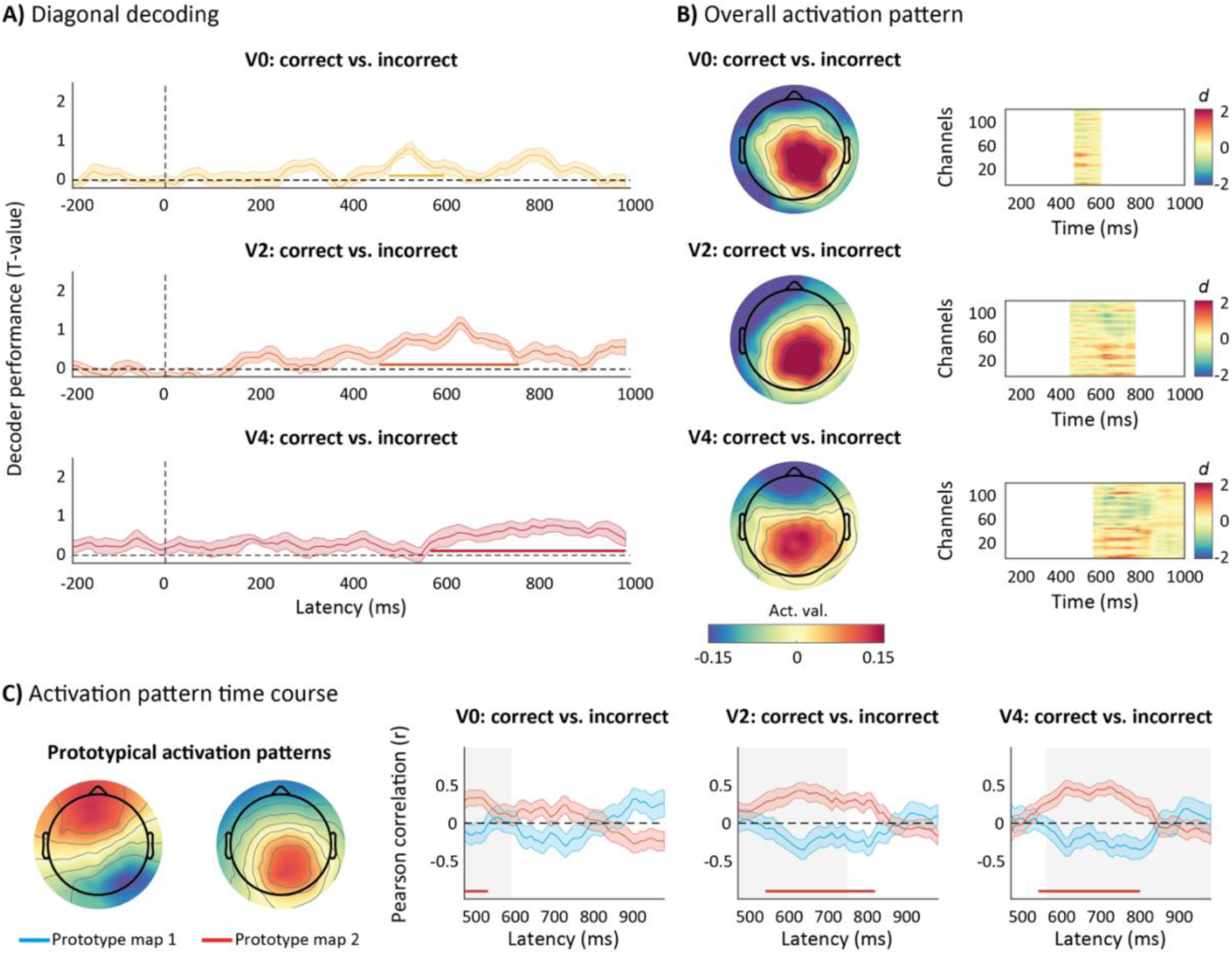
LDA between correct and incorrect reports in conditions with one vernier. A) Diagonal elements of the temporal generalization matrices (shown in Supplementary Figure 2), representing decoding results when training and testing at the same time point (group average AUC and SEM). Significant time windows are highlighted by the horizontal lines at the bottom, showing that classifiers successfully discriminate correct from incorrect offset reports in all V conditions (V0, V2 or V4; AUC > 0.5, one-tailed cluster-based permutation test, *p* < .05). B) The decoder topographies, averaged across participants, are derived from the averaged activation pattern over the entire significant window of each decoding analysis. These activations patterns were consistently expressed across participants, with many channels and time points showing large effect sizes (difference between each activation pattern and zero, calculated for each electrode and time point |Cohen’s d| > 1; bottom row). C) Prototypical activation patterns (left side) reflecting the occipital (map 1) and parietal topographies (map 2), corresponding to the average of the two distinct activation patterns identified via dissimilarity matrix analysis for all V or V-AV vs. NV contrasts (see Figures 3B and 4D), were temporally correlated with the activation patterns found in each decoding window (right side). Gray areas highlight the significant decoding window found in 6A. Blue and red lines represent the occipital and parietal topographies, respectively, and the shaded areas indicate SEM. Significant positive correlations are also highlighted in blue for the occipital topography or in red for the parietal one (Pearson’s *r* > 0, one-tailed cluster-based permutation test, *p* < .05).

In addition, the topographies obtained by averaging the main activation patterns across the entire significant decoding window (Figure 6B) were clearly distinct from those identified in the previous decoding analyses (V0 vs. V-AV and V0-AV2 vs. V0-AV4; Figure 5B). Correlations with prototypical activation patterns confirmed that decoding of the correctly reported direction was driven by a single EEG activation pattern (Figure 6C), which in this case aligned with the parietal topography identified when decoding conditions with one or two offsets from trials without any offset (Figures 3B and 4D). This topography exhibited latency shifts as a function of the vernier’s temporal position within the stream and appeared to persist longer when the vernier occurred later in time (see also Supplementary Figure 3 for analogous effects in V–AV conditions, depending on whether the first or second vernier dominated the integrated percept).

Thus, the later decoding of correct versus incorrect responses—compared to the earlier decoding reflecting differences in the spatiotemporal location or number of verniers in the stream—indicates that this centro-parietal topography reflects a neural correlate of a later stage, involving processes associated with the consciously perceived vernier offset, rather than the initial stage of unconscious integration.

## Discussion

Previous studies have proposed that in the SQM, and in related postdictive phenomena, conscious perception is preceded by a period of unconscious processing (Choi & Scholl, 2006; Wu et al., 2009; Elliott & Giersch, 2016; Sergent, 2018; Keuninckx & Cleeremans, 2021; Herzog et al., 2016, 2020). In the SQM, a sequence of lines and vernier offsets is unconsciously integrated for up to 450 milliseconds before a conscious percept emerges (Drissi-Daoudi et al., 2019; Vogelsang et al., 2023). Here, we used time-resolved decoding analysis to identify EEG markers of this unconscious integration stage and to track the transition toward the final integrated percept.

We found that each vernier offset in the stream could be decoded from EEG activity time-locked to the moment it was shown, with decoding emerging around 200 ms later (Figure 2). This was the case even though the consciously perceived vernier offset does not depend on the number or specific spatiotemporal location of the verniers within the SQM stream. Importantly, individual verniers remained decodable even when two opposite verniers were presented and their offsets were integrated (Figure 4). These results indicate that, even when individual vernier offsets are not consciously perceived, their presence and temporal position can still be inferred from neural activity.

The decoding of individual vernier offsets was driven predominantly by occipital activity (see Figures 3 and 4), with a topography resembling the N170/VAN ERP component (see also Supplementary Figure 4 for complementary ERP analyses). Although this component has often been proposed as an early correlate of conscious perception (Ojanen et al., 2003; Koivisto & Revonsuo, 2010; Railo et al., 2011; Dembski et al., 2021), several studies have related it to response to unseen stimuli (Sterzer et al., 2009; Harris et al., 2013; Cohen et al., 2024). Our results suggest that the N170/VAN may reflect an unconscious rather than a conscious processing stage, consistent with studies linking this component to the unconscious accumulation and integration of visual information (Fahrenfort et al., 2017; Noorman et al., 2025).

The N170/VAN-like component was also observed when decoding between conditions involving one or two verniers (Figure 5), even though both conditions produced the same integrated percept, namely, a single vernier offset perceived throughout the stream. Because these conditions differed in the location or number of physical offsets, this finding suggests that the topography reflects differences in the sequence of unconsciously processed events, supporting its role as a neural correlate of unconscious integration. It is also worth noting that this topography was right-lateralized (ipsilateral to the stimulus). The origin of this lateralization remains unclear, and future studies should assess the effect of attending to the left versus the right stream, as participants in the present study attended only to the right stream. One possibility is that the right hemisphere dominates processing of motion-related signals from both visual hemifields (Chiappini et al., 2022; Pitcher, 2022), or, more generally, that it reflects a right-hemisphere specialization for visuo-spatial processing (Corballis, 2003).

Following this occipital topography, we identified a distinct parietal topography (see Figures 3 and 4). This parietal topography contributed to decoding conditions with one or more verniers against the condition with no vernier, but it did not emerge when decoding between conditions containing one versus two verniers. This suggests that it carries information about the integrated percept, regardless of whether it arises from a single vernier or from the integration of two. Consistent with this interpretation, a similar centro-parietal topography was observed when decoding correct versus incorrect behavioral reports of the offset in conditions with a single vernier (see Figure 6). This topography emerged after approximately 450 ms and persisted for several hundred milliseconds, resembling the P300 ERP component (see also Supplementary Figure 4 for complementary ERP analyses).

The P300 is often described as a neural correlate of conscious access (Sergent et al., 2005; Del Cul et al., 2007; Kouider et al., 2013; Weaver et al., 2019; Ye et al., 2019; Noorman et al., 2025) and has been linked to late-access models of consciousness, where conscious perception arises from higher-order, distributed or recurrent neural processes (Baars, 2002; Dehaene et al., 2006; Lau, 2007; Dehaene & Changeux, 2011; Naccache, 2018; Seth & Bayne, 2022). However, recent studies suggest that, rather than reflecting the neural correlates of conscious perception, the P300-like component may index post-perceptual processing stages, such as decision-making and response-related activity, because it is often absent in no-report paradigms or for task-irrelevant but consciously perceived stimuli (Pitts et al., 2014; Schlossmacher et al., 2020; Cohen et al., 2020; Sergent et al., 2021). In our task, a direct motor contribution can be ruled out, as responses occurred much later, around 900 ms. Nevertheless, a post-perceptual interpretation remains possible, and future studies incorporating no-report paradigms will be important to further clarify the role of this component.

Here, we suggest that the critical shift from unconscious processing to the unified integrated percept is not reflected in the P300-like component itself, but rather in the transition from the occipital to the parietal topography. This interpretation is further supported by the close temporal alignment between this transition (Figures 3 and 4) and the window of unconscious integration previously estimated from behavioral measures (∼290-450 ms; Drissi-Daoudi et al., 2019; Vogelsang et al., 2023).

Thus, our results support the existence of long-lasting unconscious processing windows, which have been reported in other postdictive paradigms beyond the SQM (Sergent et al., 2013; Akyürek & Wolff, 2016; Thibault et al., 2016; Xia et al., 2016; Sun et al., 2017), and even with briefly presented stimuli. For example, when a 30 ms vernier is followed by a 30 ms anti-vernier, participants perceive a single integrated vernier without any percept of apparent motion (Scharnowski et al., 2009). Yet, transcranial magnetic stimulation (TMS) can alter this integrated percept up to 360 ms after stimulus onset, shifting dominance toward one of the two initial verniers depending on the timing of the TMS pulse (Scharnowski et al., 2009). These findings collectively support the idea that unconscious integration can persist well beyond the physical presence of the stimulus (see also Pilz et al., 2013 for similar results with other visual features than vernier offsets).

Although the unconscious integration window appears relatively long in the SQM, consistent with the fact that individual vernier offsets are not perceived as separate events, its duration may also reflect the demands of motion processing, whereas in other tasks shorter integration windows may allow conscious perception to emerge earlier. Indeed, our results show that the temporal dynamics of unconscious and conscious processing stages are not fixed and may vary even within the same paradigm. For example, when decoding conditions with a single vernier against conditions without any vernier (Figures 2 and 3), the latency between the vernier presentation and the onset of significant decoding decreased when the vernier appeared later in the stream (from 240 ms in V0 vs NV to 170 ms in V4 vs NV; Figure 2B). In these conditions, the occipital topography persisted for a longer duration, whereas the parietal one was shortened (Figure 3A). Moreover, the onset and duration of the second topography appeared to depend on the spatiotemporal location of the vernier dominating the unconscious integration process (Figure 6A; Supplementary Figure 3). One possibility is that the occurrence of a vernier offset initiates an unconscious processing window, while subsequent offsets or the termination of the SQM stream may accelerate the collapsing of the sequence into a unified conscious percept (Wolff & Akyürek, 2026). Although the exact nature of these non-linearities remains unclear, our results indicate that the temporal dynamics of unconscious and conscious processing stages are flexible and may be influenced by subtle details of the stimulus sequence, in contrast to standard ERP analyses, which often treat processing latencies as fixed.

In general, the fact that neural activity is time-locked to a stimulus (see Figure 2) does not imply that conscious perception occurs at that moment (the ‘vehicle problem’, Dennett & Kinsbourne, 1992; Herzog et al., 2020; Dainton, 2023; White, 2023), just as a neuron that encodes “red” is not itself red. This contrasts with theories assuming a close mapping between stimulus timing and the moment of perception (e.g., the ‘brain-time’ view, Holcombe, 2014) and implies that neither the transition between the two topographies nor the duration of the parietal topography necessarily reflects neural events involved in conscious perception itself. Instead, these neural signatures may act as precursors to consciousness or, as mentioned, reflect post-perceptual processes operating on the content of the percept. Nevertheless, this transition marks the shift from the unconscious encoding of the physical, retinotopic properties of the vernier offset—which are not perceived consciously—to the integrated representation of the stream, which forms the basis of conscious perception in the SQM. Thus, while the exact timing of conscious perception cannot be inferred directly from these topographies, they reveal the sequential neural processes that enable the emergence of a coherent percept.

In line with this, throughout the manuscript we use the terms conscious processing and conscious perception to refer to the stage in which an integrated percept of the entire SQM stream has emerged. These terms are used in contrast to the unconscious processing of individual offsets in the SQM, which participants cannot perceive individually. We must clarify, however, that by these terms we do not intend to imply or distinguish between phenomenal consciousness (i.e., the felt quality of a conscious mental state) and access consciousness (i.e., the functions enabled by conscious processing, such as behavioral reports), as our paradigm is not designed to address this distinction (Block, 2007; Michel & Doerig, 2022). As mentioned, we use the terms broadly to include also post-perceptual, decision-making processes associated with the emergence of an integrated percept.

The presence of two distinct topographies, and their temporal dynamics, is compatible with a recently proposed two-stage model of perception in which a prolonged period of unconscious processing precedes the emergence of a unified conscious percept (Herzog et al., 2016, 2020). In this framework, the first stage involves the precise encoding of each vernier offset and its temporal position, with this information maintained in a high-capacity buffer where integration operates unconsciously. This stage aligns with the independent neural encoding of individual verniers observed in our decoding results and reflected in the occipital topography (Figures 3, 4 and 5). In the second stage, the unconscious processing window enables a form of sense-making: the brain resolves the buffered information into the most coherent percept over the preceding hundreds of milliseconds. At the end of this stage, the buffered information collapses into a conscious percept that reflects the integrated structure of the sequence. This transition, and the emergence of a unitary percept, align with the decoding results associated with the parietal topography (Figures 3, 4 and 6). Overall, these sequential stages are consistent with discrete retentional models in which perception is momentary but temporally rich, delayed relative to stimulus onset, yet encoding quantitative details such as duration, speed, and direction (Herzog et al., 2016). Such dynamics do not contradict the existence of rapid visual detection (Thorpe et al., 1996) or the effects of masking (Breitmeyer & Öğmen, 2007). Detection, object recognition, and even action can unfold during unconscious processing without immediately giving rise to conscious awareness (Dehaene et al., 1998; Mitroff & Scholl, 2005; Van Gaal et al., 2012; but see also Biderman & Mudrik, 2018; Mudrik & Deouell, 2022).

While many previous findings support the long-lasting unconscious integration of visual features in the SQM (Otto, 2006; Otto et al., 2010; Plomp et al., 2009; Drissi-Daoudi et al., 2019; Vogelsang et al., 2023, 2024), the typical use of a forced-choice task based on the offset direction of the first vernier, as employed here, does not allow one to directly infer the exact percept experienced by the participants in the V–AV condition. That is, one cannot infer whether participants perceive a perfectly straight line or a weaker residual offset resulting from non-uniform integration. A recent study showed that the reported offset direction in V–AV conditions, whether consistent with the first or the second vernier, can be predicted by pre-stimulus EEG fluctuations in the alpha band (Menétrey et al., 2023). This finding suggests that integration may indeed vary across trials, with fluctuations in the relative dominance of the first or second vernier. Importantly, regardless of whether integration is uniform or non-uniform, our results show that the spatiotemporal location of each individual vernier, although not consciously accessible as a separate event because all elements are integrated within the SQM stream, can nevertheless be decoded from a specific occipital topography. Considering the role of pre-stimulus oscillatory activity, future studies could further investigate the contribution of specific frequency bands to the unconscious and conscious stages involved in SQM processing, as the present analyses focused exclusively on ERP-based decoding.

In sum, our findings support the presence of sequential processing stages in the SQM and related phenomena, with an extended period of unconscious visual processing followed by a transition to a unified, conscious percept. This has direct relevance for efforts to identify the neural correlates of conscious perception, as most theoretical frameworks, except for the global neuronal workspace theory (Dehaene et al., 2006; Dehaene & Changeux, 2011), do not explicitly separate unconscious from conscious processing stages (for reviews, see Aru et al., 2012; Storm et al., 2017). Acknowledging this two-stage structure may therefore provide a useful foundation for future research on the temporal organization of conscious and unconscious visual processing.

## Materials and Methods

### Participants

A total of 18 naive, healthy participants (9 females; age range: 18-23 years old) were recruited for the experiment. Participants had normal or corrected to normal vision, as assessed through the Freiburg acuity test (threshold for inclusion: >1; Bach, 1996). All participants signed informed consent before the experiment and received monetary compensation upon its completion. The experiment was conducted with the approval of the local ethics commission (Commission cantonale d’éthique de la recherche sur l’être humain, Canton of Vaud, Switzerland; protocol number: 2021-02270; title: Fundamental aspects of object recognition) and in accordance with the Declaration of Helsinki (except for pre-registration).

### Apparatus

The stimuli were presented on an ASUS VG248QE LCD monitor with a resolution of 1920 × 1080 pixels, a screen size of 24.5 inches, and a refresh rate of 144 Hz. MATLAB R2022b (MathWorks Inc., Natick, MA, USA) along with Psychtoolbox (Brainard, 1997) was used to generate the stimuli. During the experiment, participants were seated at a distance of 1.5 meters from the screen in a dimly lit room. The stimuli appeared as white with a luminance of 100 cd/m², displayed against a uniform black background with a luminance of 1 cd/m².

### Stimuli and experimental procedure

We used the sequential metacontrast paradigm (SQM), introduced in Otto et al. (2006), which creates the perception of two motion streams diverging from the center (Figures 1A and 1B). The sequence always began with the presentation of a central vertical line, comprising an upper and a lower segment (the length of each segment is 26.6 arcmin, with a width of 1.2 arcmin) separated by a small vertical gap (2.5 arcmin). Next, pairs of flanking lines appeared progressively further away from the center (horizontal separation between each line is 5 arcmin). The central line and all subsequent pairs of lines were presented for 27.8 ms each, with an interstimulus interval (ISI) of 20.8 ms separating each presentation. The entire stimulus sequence consisted of one central and 5 flanking lines, lasting a total duration of 270.8 ms (Figure 1A).

Participants were instructed to always covertly attend to the right stream. In a randomized order, 6 different conditions were presented (Figure 1C): In the *no vernier* control condition (NV), all lines were straight. In three experimental conditions, one line of the attended stream was offset, with the lower segment shifted either towards the right or the left relative to the upper segment (referred to as a *vernier offset* or V). In these conditions, the vernier offset was presented either in the first central line (V0 condition), in the second flanker line (V2 condition; 100 ms after stimulus onset) or in the fourth flanker line (V4 condition; 200 ms after stimulus onset). In the two last conditions, the central line always had a vernier offset, while another line in the right stream was offset in the opposite direction (referred to as an *anti-vernier* or AV). In these conditions, the opposite vernier was presented either in the second flanker line (V0-AV2 condition) or in the fourth flanker line (V0-AV4 condition). The direction of the vernier offsets was randomly determined before each trial.

In all conditions, participants were asked to indicate whether they perceived a left or right offset along the stream (Figure 1B), even when no vernier was presented. Participants were informed that two opposite verniers could also be presented in the stream and were instructed to report the first one in case they perceived both.

Participants completed a total of 12 blocks, with each block consisting of 96 trials (192 trials per condition, 1152 trials in total). Before each trial, a central fixation point was presented for 250 ms, followed by an inter-stimulus interval of 750 ms. After the SQM presentation, participants had 3 seconds to give their response by clicking one of two hand-held buttons. If no response was made within this time, the trial was discarded and repeated at the end of the block. In order to eliminate potential early responses that occurred before the complete presentation of the stimulus sequence, we also excluded trials with reaction times below 300 ms after SQM onset (average number of discarded trials: 8.3 ± 21.2). The inter-trial interval was set at 1 second.

### Offset calibration

Before the experiment proper, the offset sizes (i.e., the horizontal displacement between their upper and lower segments) were determined to achieve comparable performance levels across participants. Streams of straight lines with one single vernier were presented, and a parameter estimation by sequential testing (PEST) procedure (Taylor & Creelman, 1967) was used to adaptively determine offset sizes leading to around 70% to 80% performance. This was performed with a vernier offset separately at the three different spatiotemporal locations tested in the main experiment: the central line (V0), the second flanker line (V2), or the fourth flanker line (V4).

### Behavioral analysis

Performance was determined as the proportion of responses that matched the direction of the first presented vernier offset (Figure 1D). We compared performance between the NV condition and each V or V-AV conditions by means of paired t-tests. The NV condition served as a baseline condition; in this condition performance was determined by comparing the response (left or right reported offset) to a randomly chosen notional offset (mean accuracy was 49% ± 4%). All statistical comparisons were Bonferroni-Holm corrected for multiple comparisons (bonf_holm()). When the vernier is presented without an anti-vernier, vernier discrimination should be approximately 75%, in line with the calibration performance. Conversely, when an anti-vernier follows the first (central) vernier, the discrimination of this vernier offset should be around 50%, given the integration of the two verniers. Additionally, we anticipated no performance differences related to the specific spatiotemporal locations of the verniers and anti-verniers, as they fell within the same integration window (Drissi-Daoudi et al., 2019). Behavioral performance aligned with this expectation (see Results section).

Across the six tested conditions, average reaction times ranged from 885 ms to 920 ms, with no statistically significant difference between conditions (one-way ANOVA, F(5, 102) = 0.08, *p* = .99).

### EEG recordings and preprocessing

The EEG data were recorded using a Biosemi Active Two system (Biosemi, Amsterdam, the Netherlands) with a total of 128 Ag–AgCl active electrodes, providing comprehensive scalp coverage. The positioning of the cap was adjusted individually to ensure the Cz electrode was equidistant from the inion and nasion, as well as equidistant from each ear. In addition, the electrooculogram (EOG) was recorded with 4 electrodes positioned 1 cm above and below the right eye and 1 cm lateral to the outer canthi. The recording was referenced to the CMS-DRL ground, maintaining the montage potential close to amplifier zero through a feedback loop. The sampling rate during recording was set at 2048 Hz.

For EEG preprocessing, EEGLAB was utilized (version v2021.1; Delorme et al., 2011; Delorme & Makeig, 2004). The EEG data were first downsampled to 250 Hz. To remove linear trends, detrending was applied (de Cheveigné & Arzounian, 2018), followed by a lowpass filter with a cutoff frequency of 40 Hz. The EEG data were then epoched from −1 s to 1.5 s relative to the onset of the SQM. To ensure data quality, a visual inspection was conducted to identify and exclude epochs (pre-selected with *pop_jointprob* function) or channels with significant noise or artifacts. Epochs containing eye-related artifacts (e.g., blinks or saccades) overlapping with, or occurring around, the SQM presentation were also removed (but see also Drissi-Daoudi et al., 2020 for evidence that integration between two verniers is preserved even when they are presented before and after an eye movement). Then, an ICA decomposition (*pop_runica* function with Picard algorithm; Ablin et al., 2017a, 2017b) was performed, with a temporary interpolation of the removed channels to maintain a consistent 128-channel configuration across all participants. The ICA decomposition also included “fake” vertical and horizontal EOG recordings, computed from the 4 EOG channels (differences between the channels located above and below the right eye, and between the channels located on the lateral side of each eye). The visually identified independent components associated with eye or muscular artifacts (pre-selected with *pop_icflag* function) were subsequently removed. Lastly, a final interpolation of the removed channels was conducted, along with an average reference including only the EEG channels.

In total, 6.8% of the electrodes were interpolated, while 3.2% of the epochs and 9.1% of the independent components were removed during the preprocessing procedure.

### EEG analysis

We aimed to investigate whether brain activity reflects unconscious or conscious processing stages by decoding differences in EEG dynamics in two configurations of the SQM: 1) when only a single vernier is presented at different spatiotemporal locations within the stream (V conditions), with all these conditions resulting in the same percept (i.e., one perceived vernier); and 2) when two opposing verniers are presented (V-AV conditions), yet there is no conscious percept of the individual offsets as they integrate.

### EEG decoding

Linear discriminant analysis (LDA) with temporal or cross-condition generalization was used to investigate neural representations related to different SQM conditions or percepts. We implemented LDA with custom-made functions written in MATLAB, based on the recommended settings for EEG data (Subasi & Ismail Gursoy, 2010; Grootswagers et al., 2017). For each participant, LDA with temporal generalization was conducted with a leave-one-pseudo-trial-out cross-validation routine (500 iterations). In each iteration, 80% of the trials were sampled and combined into pseudo-trials (average of 20 trials) for the two classes, i.e., the SQM conditions, that were decoded. The mean of the training set was removed to both testing and training sets, and classifier weights were estimated using a regularized covariance (Ledoit & Wolf, 2004; Haufe et al., 2014; Kayser et al., 2016). The classifier weights, obtained from the training set for each time point, were applied to predict the classes in the EEG data of the testing set. Performance was evaluated using the area under the curve (AUC). This process was repeated for each time point (cross-temporal decoding, i.e., each classifier is tested on all time points), ranging from −200 to 1000 ms, with a sliding window of 7 samples corresponding to a resolution of 70 ms (Grootswagers et al., 2017). LDA with cross-condition generalization was performed similarly and with identical parameters, except that the classes used in the training sets differed from those used in the testing sets. For all decoding analyses, EEG data were resampled at 100Hz and z-scored.

We started by examining V conditions, where the presentation of a single vernier in the stream results in a clear perception of an offset. We used LDA with temporal generalization to discriminate between trials from the control condition (NV) and trials from conditions presenting one single vernier (V0, V2, or V4), irrespective of the behavioral responses (i.e., correct or incorrect report of the vernier offset; Figures 2A and 2B). We also evaluated cross-condition generalization between V conditions. Using classifiers trained with V2 or V4 vs. NV conditions, we decoded V0 vs. NV conditions (Figure 2C, see also Supplementary Figure 1 for additional decoding analyses between V conditions).

Second, we investigated the neural representations in the V-AV conditions, where the two opposite verniers integrate with each other. LDA with temporal generalization was used to classify between trials from the NV condition and one of the two conditions presenting two opposite verniers (V0-AV2 or V0-AV4), irrespective of the behavioral responses (i.e., correct or incorrect report of the first vernier offset; Figure 4A; see also Figure 4F for cross-condition generalization between V and V-AV conditions and Supplementary Figure 3 for additional cross-condition generalization between V and V-AV conditions, as a function of whether the first or second vernier dominated the reported percept). Next, we conducted additional decoding analyses between V0 and V0-AV2 or V0-AV4 conditions, as well as between the two V-AV conditions (Figure 5).

Lastly, we conducted an LDA analysis with temporal generalization to decode between correct and incorrect reports of the offset within each condition presenting one vernier (V0, V2, and V4; Figure 6). This analysis aimed to specifically identify neural correlates that reflect conscious processing. All parameters remained consistent with those listed above, except for the averaging of trials for pseudo-trials, which was adjusted to 4 due to the reduced number of available trials, especially the incorrect trials (comprising only ∼30% of the total for each condition).

### Statistical evaluation of decoding analyses

For all decoding analyses, either with temporal or cross-condition generalization, the statistical evaluation of the 2D matrices of cross-temporal decoding results (participants x time x time) employed cluster-based permutation approaches and surrogate analysis, following established methodologies (Nichols & Holmes, 2002; Maris & Oostenveld, 2007; Kayser et al., 2016). Clusters were defined as consecutive time points where the decoding successfully exceeded chance levels (chance = 0.5, paired t-test with α = 0.05). The cumulative t-values within each cluster were then compared to the maximum sum derived from surrogate clusters (permutations = 10’000). Time points were considered statistically significant if the probability in the surrogate data was <0.05 for the corresponding clusters.

### EEG activation patterns

We estimated activation patterns from the LDA classifiers using the regression-based approach (eq. [8] in Haufe et al., 2014). Briefly, this method applies an ordinary least squares regression to derive a pattern that, when multiplied by the discriminant signal obtained via LDA, best approximates the original EEG data used as input to the decoder (electrode x time points x pseudotrials). This approach yields a ‘forward model’ that allows for neurophysiologically interpretable patterns, addressing the known issue of non-interpretable decoder weights (Haufe et al., 2014; Grootswagers et al., 2017).

We assessed the temporal dynamics of the activation patterns in the context of each decoding analysis (V0, V2, or V4 vs. NV, Figure 2; V0-AV2 or V0-AV4 vs. NV, Figure 4; V0-AV2 or V0-AV4 vs. V0, V0-AV2 vs. V0-AV4, Figure 5; correct vs. incorrect reports in V0, V2, and V4, Figure 6; see also Supplementary Figure 1 for V0 vs. V2 or V4, and V2 vs. V4). For each participant, we extracted the activation patterns over the significant time points.

Across the five initial decoding contrasts (i.e., all V vs. NV and all V-AV vs. NV conditions; see Figures 2 and 4), we first computed time-by-time dissimilarity matrices based on the spatial topographies of decoder activation in order to characterize the temporal structure of these EEG activation patterns. These activation patterns were then averaged across participants, and the resulting maps were z-scored across electrodes at each time point to control for overall amplitude differences. Next, we calculated pairwise dissimilarities between all time points within the significant decoding window using the Euclidean distance—a standard measure of topographic dissimilarity that quantifies how different two z-scored maps are by summing the squared differences across electrodes.

This procedure yielded a symmetric dissimilarity matrix that reflects the similarity structure of spatial patterns over time. To identify periods during which activation patterns remained relatively stable (i.e., dominated by similar topographies), we transformed this dissimilarity matrix into a graph representation, where each node corresponded to a time point and edges were defined between pairs of time points with dissimilarity values below the median of the entire matrix. This thresholding ensured that only relatively similar patterns were considered connected in the graph.

We then applied the Louvain community detection algorithm (Blondel et al., 2008; Rubinov & Sporns, 2010), a widely used method for detecting modular structure in networks, to this graph. In this context, the algorithm grouped together contiguous time points that shared similar activation topographies, thereby identifying clusters of temporally stable EEG patterns. To mitigate potential instabilities in the clustering due to noise at the single-participant level, this procedure was performed on group-averaged data. Nevertheless, we verified the robustness of the main findings across participants by estimating effect sizes (see below). The data-driven clustering allowed us to define distinct temporal clusters characterized by dominant topographic maps (Figures 3A and 4C), which were used in subsequent analyses.

Topographies of the main activation patterns were then generated by averaging over temporal windows corresponding to the identified clusters (Figures 3B and 4D), revealing two highly consistent spatial topographies. To examine their temporal characteristics and contributions to the overall dynamics, we correlated these spatial patterns with the temporal evolution of EEG activation (back-projected time courses; Figures 3C and 4D). For each time course independently, clusters were defined as consecutive time points where the correlation significantly exceeded 0 (paired t-test with α = 0.05). The cumulative t-values within each cluster were then compared to the maximum sum derived from surrogate clusters (permutations = 10’000). Time points were considered statistically significant if the probability in the surrogate data was < 0.05 for the corresponding clusters. Notably, while the topographies derived from dissimilarity matrices were based on group-averaged data, the back-projection plots demonstrate the consistency of these maps and their temporal dynamics across individuals. To further assess these effects at the individual level, we quantified the proportion of participants whose activation patterns showed positive correlations with the dominant topographic maps within the time windows corresponding to the two temporal clusters defined by these maps (Pearson’s r > 0 when averaged across each respective time window). It should also be noted that, because the two topographies exhibited largely opposing dipolar scalp distributions, their back-projection onto individual EEG activation patterns resulted in anticorrelated time courses. Interpretation should therefore rely primarily on periods of positive correlation, as these indicate time windows in which the corresponding topography was consistently expressed across participants.

To further assess the reliability of the distinct maps across participants, we estimated Cohen’s *d* (abbreviated as d in the figures) at each electrode and time point by comparing the corresponding activation values across participants against zero (Figures 3A and 4C). The effect sizes here quantify the extent to which activation at a given electrode and time point was consistently different from zero across participants. The presence of large positive and negative effect sizes therefore indicated that the observed topographic patterns were robust and reliably expressed across individuals.

For the additional decoding contrasts (i.e., V0 vs. V-AV conditions, V0-AV2 vs. V0-AV4 and correct vs. incorrect reports in V conditions; see Figures 5 and 6; see also Supplementary Figure 1 for V0 vs. V2 or V4, and V2 vs. V4), topographies of the main activation patterns were generated by averaging over the entire significant decoding window (Figures 5B and 6B). In addition, we averaged the two maps identified across all decoding results between V or V-AV vs. NV conditions (Figures 3B and 4D) to define two ‘prototypical’ activation patterns associated with the two distinct processing stages (Figures 5C and 6C). We then fitted these prototypes to the individual activation patterns from the additional decoding analyses using separate linear models to assess their relative contributions. For each time course independently, clusters were defined as consecutive time points where the correlation significantly exceeded 0 (paired t-test with α = 0.05). The cumulative t-values within each cluster were then compared to the maximum sum derived from surrogate clusters (permutations = 10’000). Time points were considered statistically significant if the probability in the surrogate data was <0.05 for the corresponding clusters. This analysis confirmed that each decoding result was dominated by a single topography (Figures 5C and 6C). This was further complemented by the estimation of Cohen’s *d* for the difference between each activation pattern and zero at the level of individual electrodes and time points, which highlighted a single, consistent pattern across each significant time window (Figures 5B and 6B).

## Acknowledgments

We sincerely thank Marc Repnow for his excellent technical support. We are also grateful to the Reviewers for their valuable comments and suggestions, which have greatly improved the quality of our manuscript.

## Data Availability

The data supporting the findings of this study are available on the Open Science Framework (https://osf.io/d83vs/)

## Code Availability

All scripts necessary to reproduce the analyses and the figures are available on GitHub (https://github.com/MaelanMenetrey/EEGdecoding_SQM).

## Authors contribution statements

M.Q.M, M.H.H. and D.P. conceived the study; M.Q.M collected the data; M.Q.M and D.P. analyzed the data; M.Q.M, M.H.H. and D.P. interpreted the results; M.Q.M, M.H.H., and D.P. wrote the manuscript.

## Supplementary Material

### Decoding analyses between V conditions

As supplementary analyses, we used linear discriminant analysis (LDA, see Methods) to discriminate between conditions presenting a single vernier (V0, V2, or V4), irrespective of behavioral responses (i.e., whether participants correctly or incorrectly reported the vernier offset). Although these conditions differed physically, because the single vernier appeared at different spatiotemporal locations, they did not differ perceptually (see Otto, 2006; Drissi-Daoudi et al., 2019): in all cases, the offset is integrated into the motion stream and can be reported. The aim was therefore to confirm that the occipital topography, which we associated with unconscious processing of the individual offset, contributed most strongly to decoding performance.

The classifiers (V0 vs. V2 or V4, and V2 vs. V4) successfully discriminated between all pairs of conditions (Area Under the Curve, AUC > 0.5, one-tailed cluster-based permutation test, *p* < .05; Supplementary Figure 1A). In addition, the onset and duration of significant decoding windows varied systematically depending on the conditions being compared. When V0 was included in the comparison, decoding began around 200 ms, similar to previous analyses comparing V0 vs. NV (Figure 2) and V-AV conditions vs. NV (Figure 4). By contrast, decoding onset was delayed to around 300 ms when comparing V2 and V4, resembling the decoding latencies observed for V2 vs. NV. Finally, when V4 was one of the compared conditions, decoding windows lasted longer, extending beyond 800 ms.

Moreover, as expected, these additional analyses confirmed the dominant contribution of the occipital topography within the significant decoding windows (Supplementary Figure 1C), indicating that decoding was driven primarily by differences in the topography we associate with unconscious processing. However, we also observed smaller contributions from the parietal topography (Supplementary Figure 1C). This may reflect the fact that the decoded conditions involved similar processes occurring at temporally shifted time points, as shown by the cross-condition decoding (Figure 2C) and the clustering analysis (Figure 3A) with the V conditions vs. NV. Consequently, the EEG activation patterns underlying decoding may represent a mixture of occipital and parietal topographies across time (Supplementary Figure 1B). Alternatively, the later dominance of parietal topography observed in comparisons involving V4 (Supplementary Figure 1C) may indicate that the temporal dynamics of parietal activity vary across conditions, potentially persisting longer in the V4 condition, in line with non-linear temporal dynamics observed across other analyses (see Discussion).

**Supplementary Figure 1.**
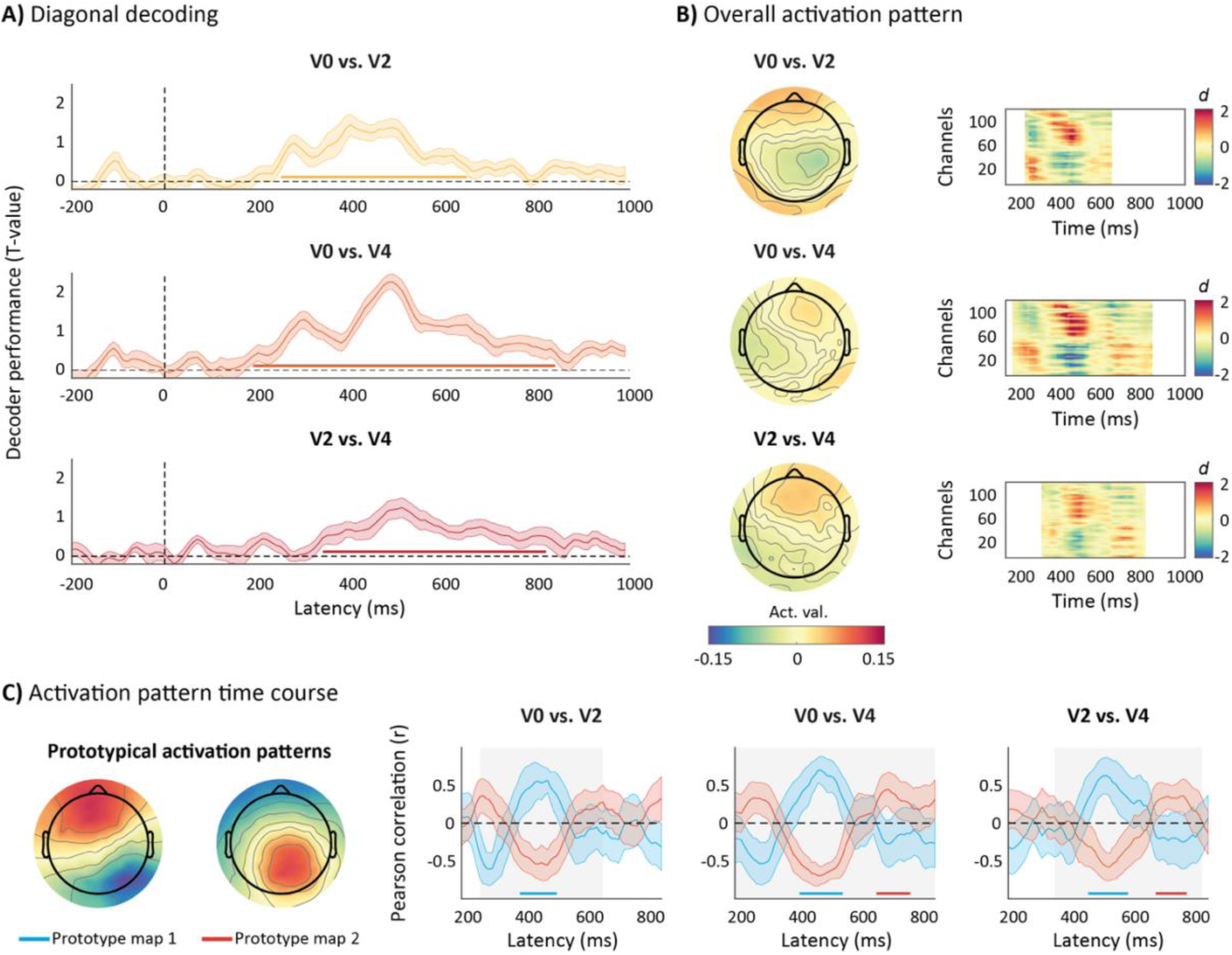
LDA between conditions with one vernier. A) Diagonal elements of the temporal generalization matrices (shown in Supplementary Figure 2), representing decoding results when training and testing at the same time point (group average AUC and SEM). Significant time windows are highlighted by the horizontal lines at the bottom, showing that classifiers successfully discriminate the conditions (V0 vs. V2 or V4, and V2 vs. V4; AUC > 0.5, one-tailed cluster-based permutation test, *p* < .05). B) The decoder topographies, averaged across participants, are derived from the averaged activation pattern over the entire significant window of each decoding analysis (left side). These activations patterns were consistently expressed across participants, with many channels and time points showing large effect sizes (difference between each activation pattern and zero, calculated for each electrode and time point |Cohen’s d| > 1; right side). C) Prototypical activation patterns (left side) reflecting the occipital (map 1) and parietal topographies (map 2), corresponding to the average of the two distinct activation patterns identified via dissimilarity matrix analysis for all V or V-AV vs. NV contrasts (see Figures 3B and 4D), were temporally correlated with the activation patterns found in each decoding window (right side). Gray areas highlight the significant window found in Supplementary Figure 1A. Blue and red lines represent the occipital and parietal topographies, respectively, and the shaded areas indicate SEM. Significant positive correlations are also highlighted in blue for the occipital topography or in red for the parietal topography (Pearson’s *r* > 0, one-tailed cluster-based permutation test, *p* < .05).

### Temporal generalization of decoding analyses

LDA with temporal generalization was used to test decoding performance across the different SQM conditions. By training a classifier at each time point and testing it across all other time points, we obtained 2D matrices of cross-temporal decoding results (participants × time × time). Cluster-based permutation tests and surrogate analyses were then used to identify significant decoding clusters (see Methods).

These temporal generalization matrices were presented in the main manuscript for decoding analyses comparing V and V-AV conditions with NV conditions (Figures 2 and 4). However, for reasons of space, they were not shown for the analyses comparing V0 with V0-AV2 or V0-AV4 conditions, and V0-AV2 with V0-AV4 conditions (Figure 5), correct versus incorrect reports in V0, V2, and V4 conditions (Figure 6), or V0 with V2 or V4 conditions, as well as V2 with V4 conditions (Supplementary Figure 1). The corresponding temporal generalization matrices for all these analyses are presented in Supplementary Figure 2 (AUC > 0.5; one-tailed cluster-based permutation test, *p* < .05).

**Supplementary Figure 2.**
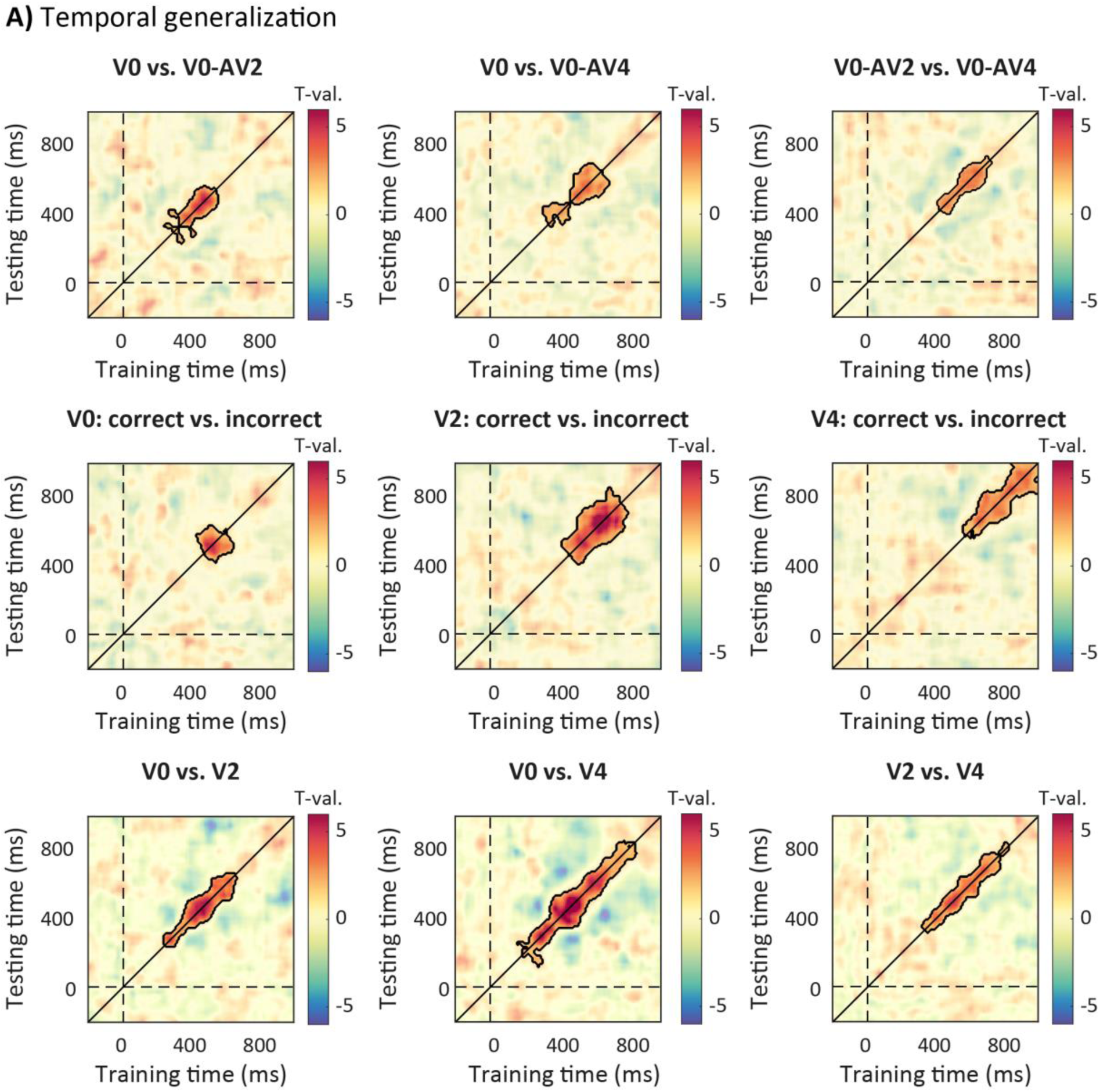
Temporal generalization matrices supporting the decoding results for the different condition comparisons shown in Figure 5 (V0 vs. V0-AV2 or V0-AV4, and V0-AV2 vs. V0-AV4), Figure 6 (correct vs. incorrect reports in V0, V2 or V4) and Supplementary Figure 1 (V0 vs. V2 or V4, and V2 vs. V4). Significant clusters are highlighted (AUC > 0.5, one-tailed cluster-based permutation test, *p* < .05).

### Temporal and cross-condition generalization as a function of the reported vernier (1^st^ or 2^nd^)

In these supplementary analyses, we decoded the V–AV conditions (V0–AV2 and V0–AV4) from the NV condition, separately for trials in which participants reported the first (i.e., V0) or the second vernier offset (i.e., V2 or V4). The aim was to further corroborate two key points: (1) that the occipital topography is time-locked to the onset of the first vernier, even in trials where the second vernier dominates the percept and report—indicating unconscious processing; and (2) that the parietal topography is time-locked to the onset of the perceived (and reported) vernier, as observed in Figure 6A.

The decoder successfully discriminated both V0–AV2 and V0–AV4 trials, regardless of the reported vernier, from NV trials (AUC > 0.5; one-tailed cluster-based permutation test, p < .05; Supplementary Figure 3A). However, the significant decoding windows showed a systematic temporal shift depending on which vernier was reported. When participants reported the first vernier, significant decoding emerged around 200 ms, similar to the analyses including all V–AV trials (vs. NV; Figure 4). In contrast, when the second vernier was reported, the decoding onset was delayed. At first glance, this pattern might appear inconsistent with the interpretation that the initial decoding and occipital topography are time-locked to the unconsciously processed vernier (which is always V0 in the V–AV conditions).

Because this analysis involved fewer trials due to the split by report, we further assessed the temporal dynamics of the two topographies by correlating the individual activation patterns with the two prototypical topographies identified in the main analyses (Figures 5C and 6C; see Methods). This correlation analysis revealed a significant match with the occipital topography time-locked to the onset of the first vernier and independent of which vernier was reported, followed by the parietal topography, which was delayed and extended in duration when participants reported the second vernier (Supplementary Figure 3B). Thus, the occipital topography—although still evident in the correlation analysis—may not have survived cluster-based permutation correction in the full temporal generalization matrices.

Together, these findings replicate and extend the main results: the occipital topography is associated with unconscious processing, time-locked to the physical onset of the first vernier (Figures 3C and 4E), whereas the parietal topography reflects the neural correlate of the consciously perceived vernier offset (Figure 6A).

In another supplementary analysis, we decoded V-AV conditions, again separately as a function of the reported vernier, from NV conditions but using classifiers trained on V conditions (V0, V2 or V4 vs. NV). Only V trials where the reported vernier offset was correct were included to ensure that any decoding results could be linked to conscious processing. This approach served two purposes:

(1) to confirm that a single vernier offset is consistently represented in V-AV conditions, even when the reported percept does not match the first offset (V0); and (2) to assess whether the temporal decoding delays previously observed in V conditions (see Figures 2B and 2C) also emerge in V-AV conditions, and whether such delays depend systematically on the spatiotemporal location of the reported vernier.

The cross-condition results successfully decoded V-AV conditions from the NV condition in all cases (AUC above 0.5, one-tailed cluster-based permutation test, *p* < .05; Supplementary Figure 3C). These results confirm that the EEG patterns related to the V-AV and NV conditions differ (see also Figure 4A) and that the neural representation of a vernier is preserved even when two opposite verniers are presented (see also Figure 4F). Moreover, they demonstrate that temporal shifts in decoding between training and testing sets, as indicated by significant clusters that are not aligned along the diagonal, directly depend on whether the first or second vernier is reported (Supplementary Figure 3C). For example, when classifiers trained on V2 or V4 were tested on the V0-AV4 condition, significant decoding clusters appeared primarily below the diagonal; Supplementary Figure 3C, lower row), indicating faster decoding in the testing set when the first (central) vernier was reported. This effect was more pronounced for classifiers trained on V4 than on V2. Conversely, when the second vernier was reported in the V0-AV4 condition, the significant decoding cluster appeared above the diagonal with classifiers trained on V2, indicating delayed decoding in the testing set. Lastly, classifiers trained on V4 led to a significant decoding cluster aligned along the diagonal, confirming the absence of temporal shift.

Together, these results confirm that the neural processes underlying V–AV conditions—regardless of which vernier is reported—closely resemble those observed in V conditions, following a similar sequence of unconscious and conscious processing stages (see Figures 2 and 3 for comparison). Moreover, consistent with the results shown in Supplementary Figure 1, they indicate that the temporal dynamics of these processes, particularly those associated with the parietal topography, vary as a function of the perceptual outcome—that is, depending on which vernier offset dominates the final percept and behavioral report.

**Supplementary Figure 3.**
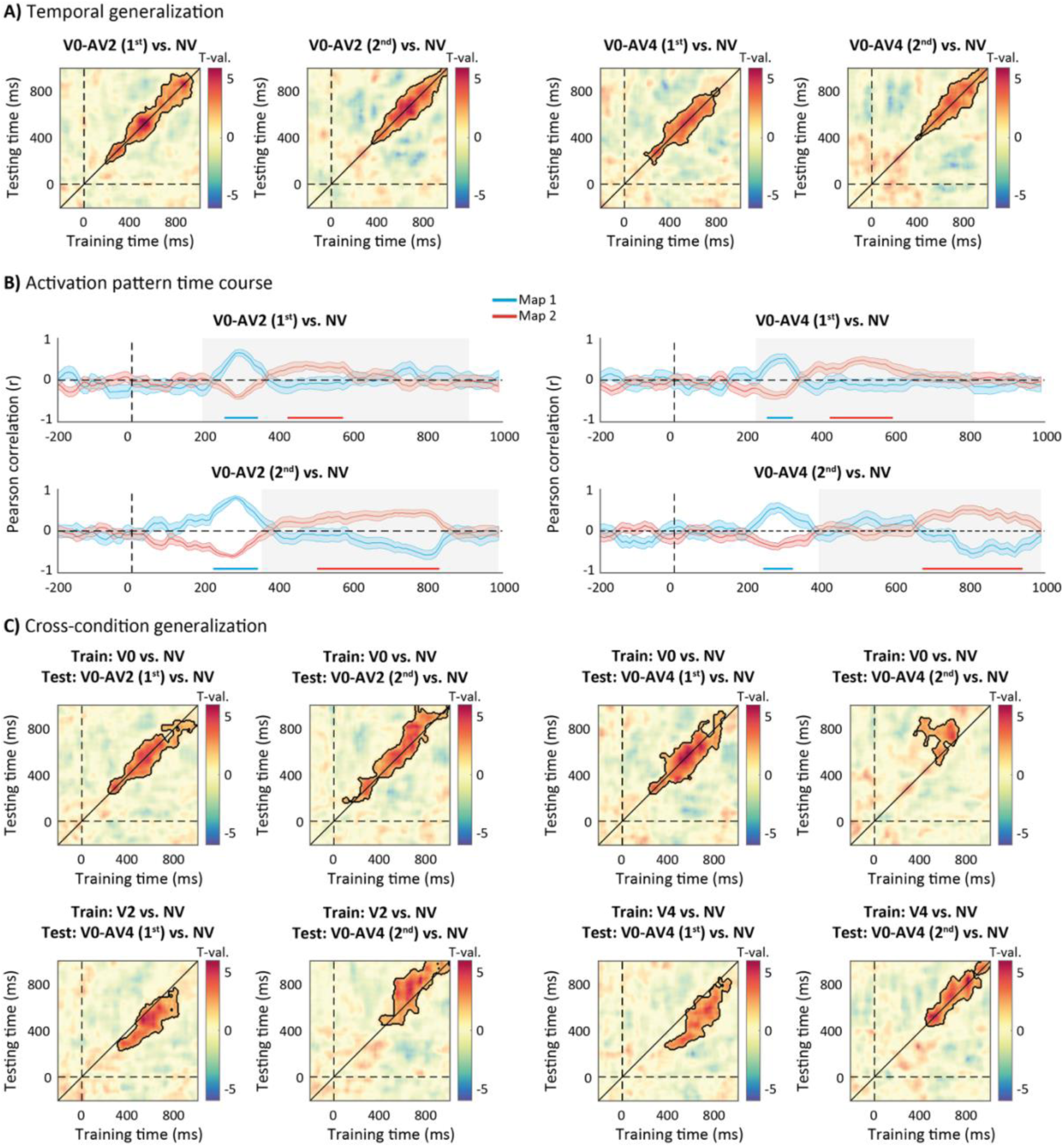
LDA with temporal and cross-condition generalization to decode conditions with two opposite verniers and no vernier. Decoding was applied separately on the V-AV trials where the first or the second vernier was reported A) Temporal generalization matrices showing that classifiers successfully discriminate the conditions (V0-AV2 or V0-AV4 vs. NV), regardless of the reported vernier (1^st^ or 2^nd^). Significant clusters are highlighted (AUC > 0.5, one-tailed cluster-based permutation test, *p* < .05). B) Prototypical activation patterns (Figures 5C and 6C), corresponding to the average of the two distinct activation patterns identified via dissimilarity matrix analysis for all V or V-AV vs. NV contrasts (Figures 3B and 4D, see Methods), were temporally correlated with the activation patterns found in each decoding window. Gray areas highlight the significant window found in Supplementary Figure 3A. Blue and red lines represent the first and second topographies, respectively, and the shaded areas indicate SEM. Significant positive correlations are also highlighted in blue for the first map or in red for the second (Pearson’s *r* > 0, one-tailed cluster-based permutation test, *p* < .05). C) Cross-condition generalization matrices showing that classifiers trained with V conditions (V0, V2 or V4) vs. NV condition successfully discriminate between the two V-AV conditions (V0-AV2 or V0-AV4 vs. NV), regardless of the reported vernier (1^st^ or 2^nd^). Significant clusters are highlighted (AUC > 0.5, one-tailed cluster-based permutation test, *p* < .05).

### Event-related potential analysis

Our decoding analyses showed that decoding performance was driven either by the presence of two successive distinct maps—namely, the occipital and parietal topography (for V or V-AV conditions vs. NV conditions; see Figures 3 and 4)—or by the presence of a single dominant map. The occipital topography contributed most strongly when unconscious processing differed despite a similar integrated percept (e.g., between V and V-AV conditions, or among V and V-AV conditions; see Figure 5 and Supplementary Figure 1), whereas the parietal topography dominated when perceptual reports differed (i.e., correct vs. incorrect reports in V conditions; see Figure 6).

In the Discussion, these two distinct topographies are compared with well-known event-related potential (ERP) components. In particular, we relate the occipital topography to the N170/VAN component and the parietal topography to the classic P300. In this additional plots, we further validate the involvement of these specific ERP components.

Condition-specific ERPs were first computed for each participant by averaging EEG epochs across trials within each condition. A pre-stimulus baseline correction was applied using the interval from −450 to −50 ms relative to stimulus onset. This resulted in a channels × time ERP matrix for each participant and condition. Overall, inspection of the grand-averaged ERPs revealed highly similar patterns across conditions (examples are shown for V0 and NV conditions in Supplementary Figure 4A).

To isolate condition-specific activity, differential ERPs were computed by subtracting the NV condition from each experimental condition (Supplementary Figure 4B). Difference maps were then normalized by the across-subject standard deviation to estimate effect sizes. The results revealed strongest effects at different latencies depending on condition, with differences emerging later in V2 and V4, consistent with the decoding onset reported in Figure 2B. Moreover, the strongest effects aligned with the two temporal windows identified by the cluster analyses (Figures 3A and 4C), and the corresponding topographies matched the EEG patterns contributing to decoding within these windows (example shown for V0 in Supplementary Figure 4C, see Figures 3B and 4D for comparison). In both windows, we identified the electrode showing maximal effect. This corresponded to an occipito-temporal electrode in the first window and a centro-parietal electrode in the second window (Supplementary Figure 4C). Because highly similar electrodes were selected across all conditions, we used these two electrodes as the basis for subsequent electrode-specific ERP plots for each condition.

For both the occipito-temporal and centro-parietal electrode, we estimated the raw ERPs (example shown for V0 in Supplementary Figures 4D and 4E) and differential ERPs (V or V-AV conditions minus NV condition; Supplementary Figures 4F and 4G). At the group level, V and V-AV conditions showed a clear increase in negative amplitude relative to the NV condition at the occipito-temporal electrode (Supplementary Figures 4F), starting around 200 ms for V0 and V-AV conditions, and emerging later for V2 and V4 conditions. For each condition, this effect closely aligned with the onset of significant decoding (vs. NV condition) and with the time window reflecting a contribution of the occipital topography to decoding (Figures 3C and 4E). These findings therefore support our interpretation that differences between V or V-AV conditions and the NV condition arise from modulation of early occipital activity, potentially consistent with a stronger N170/VAN-like response, time-locked to the onset of the first vernier in the stream. At the centro-parietal electrode, a later positive modulation distinguished V and V-AV conditions from the NV condition (Supplementary Figures 4G), with the earliest onset starting around 360 ms but varying again across conditions with later presentation of a single vernier. This pattern closely matched the second time window identified in the decoding analyses and associated with the contribution of the parietal topography (Figures 3C and 4E) and is therefore consistent with the interpretation that the later effects reflect a P300-like response.

**Supplementary Figure 4.**
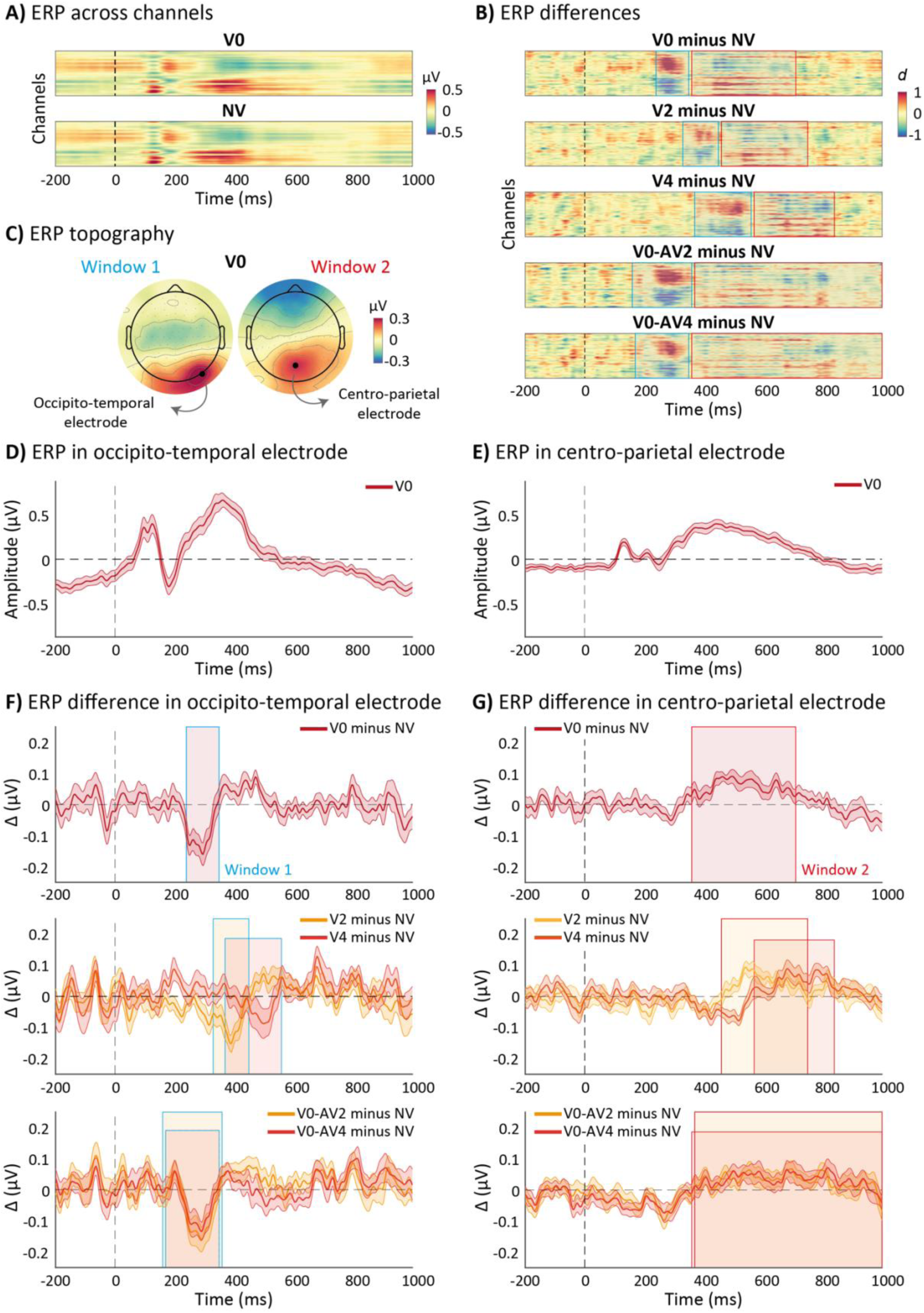
Analysis of condition-specific ERPs. A) Grand-averaged ERPs across participants and conditions at all electrodes, derived from the EEG signal. Only V0 and NV conditions are shown for illustration purposes. B) Differential ERPs obtained by subtracting the NV condition from each V and V-AV condition. Gray areas indicate the two temporal windows identified in the clustering analysis (Figures 3A and 4C). Blue and red outlines highlight the time windows associated with occipital and parietal contributions to decoding, respectively. C) ERP topographies computed by averaging ERP activity across participants within the two temporal windows identified by clustering analyses. The electrode showing maximal activity within each topography is indicated by a black dot. D–E) ERP time courses at the electrodes showing maximal activity in each time window: occipito-temporal electrode (D) and centro-parietal electrode (E). Only the V0 condition is shown for illustration. Shaded areas indicate SEM. F–G) Differential ERPs at the occipito-temporal (F) and centro-parietal (G) electrodes, computed as each condition minus NV. Colored areas indicate the two temporal windows identified by clustering analyses (Figures 3A and 4C) with blue and red outlines highlighting the time windows associated with occipital and parietal contributions to decoding, respectively. Yellow and red lines represent different V and V-AV conditions, respectively, and the shaded areas indicate SEM.

